# SpaDE: Semantic Locality Preserving Biclustering for Neuroimaging Data

**DOI:** 10.1101/2024.06.08.598092

**Authors:** Md Abdur Rahaman, Zening Fu, Armin Iraji, Vince Calhoun

## Abstract

The most discriminative and revealing patterns in the neuroimaging population are often confined to smaller subdivisions of the samples and features. Especially in neuropsychiatric conditions, symptoms are expressed within micro subgroups of individuals and may only underly a subset of neurological mechanisms. As such, running a whole-population analysis yields suboptimal outcomes leading to reduced specificity and interpretability. Biclustering is a potential solution since subject heterogeneity makes one-dimensional clustering less effective in this realm. Yet, high dimensional sparse input space and semantically incoherent grouping of attributes make post hoc analysis challenging. Therefore, we propose a deep neural network called semantic locality preserving auto decoder (SpaDE), for unsupervised feature learning and biclustering. SpaDE produces coherent subgroups of subjects and neural features preserving semantic locality and enhancing neurobiological interpretability. Also, it regularizes for sparsity to improve representation learning. We employ SpaDE on human brain connectome collected from schizophrenia (SZ) and healthy control (HC) subjects. The model outperforms several state-of-the-art biclustering methods. Our method extracts modular neural communities showing significant (HC/SZ) group differences in distinct brain networks including visual, sensorimotor, and subcortical. Moreover, these biclustered connectivity substructures exhibit substantial relations with various cognitive measures such as attention, working memory, and visual learning.

## I. Introduction

Biclustering has been successful in knowledge discovery and unveiling the latent manifold of a large dataset, especially in biological data analysis. However, the clustering quality is strongly dependent on a meaningful representation of the data. Biological data e.g., neuroimaging, and genomics often involve high dimensionality, noise, and missing values, posing challenges for a model to learn the true illustrations of data variance that lies on highly non-linear manifolds. Deep neural network (DNN) has shown impressive performance in learning cluster-friendly representation and clustering the latent space [1]. To this end, the DNN-based biclustering model is less explored, here, we propose a deep biclustering framework named semantic locality preserving auto decoder (SpaDE) for unsupervised feature learning and biclustering. Individuals with neuropsychiatric conditions like schizophrenia are heterogeneous with diverse behavioral and neurological variations [2]. Finding constrained low-level patterns is necessary in neuropsychiatric conditions to provide insights into underlying biological processes and cognitive dysfunctions instrumental for treatment and interventions [3]. However, these variations are often present in a comparatively shorter span of the data dimensions typically across a subset of individuals and features. As such, biclustering allows for stratifying both subject and feature dimensions for effective navigation through those homogeneous subgroups. Biclustering has been employed to understand brain functioning [4], temporal modulation [5], and structural changes [2].

In neuroimaging bicluster (BiC), the patterns are expected to be more disjoint, and semantically consistent. It gives more flexibility to interpret the bi-clustered community and examine the neurological relevance. Carrying similar values might not manifest the equivalent semantic meaning in biological data analysis thus SpaDE regulates semantic locality preservation (SLP) in the biclusters. SLP has been adopted in multiple research areas including genomics [6], proteomics [7], and natural languages [8]. SLP maximizes semantic consistency and aligns latent data points to their intrinsic manifold. So, the bi-clustered neural attributes are more plausible to be participating in coherent brain functionalities. Current studies show neuropsychiatric disorders as a disease of brain connectivity [9]. So, effective connectivity is one of the areas that is widely used for better understanding the interactions between brain regions [10]. Thus, our study aims to explore the influential communities in the human brain connectivity dynamic and we apply our proposed method to the static functional network connectivity (sFNC). SpaDE unveils semantically curated subgroups of connections and relates them with diverse cognitive scores for the potential explanations of cognitive deficits in schizophrenia. Moreover, the subgrouped connectivity patterns exhibit significant HC/SZ group differences (Fig. 2).

**Fig. 1.**
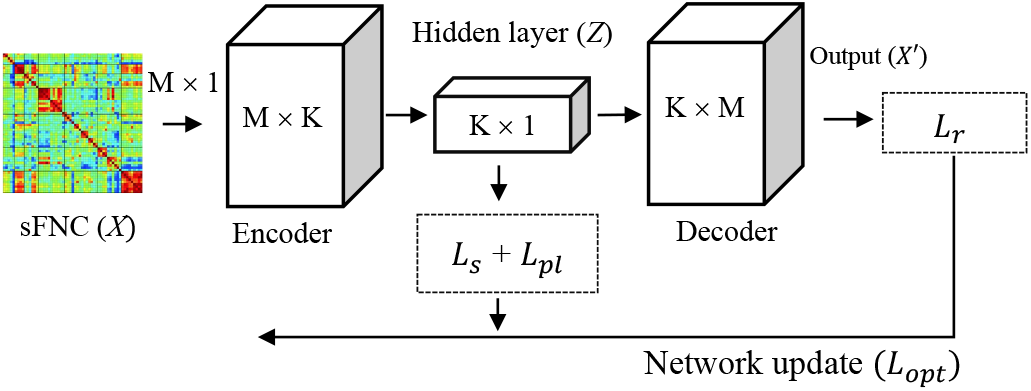
Our proposed SpaDE model - an encoder-decoder architecture. Here *L*_*r*_, *L*_*s*_, and *L*_*pl*_ represent the reconstruction loss, sparsity, and semantic locality constraints, respectively. The bottleneck layer (hidden layer) has k neurons which also illustrate the number of expected biclusters. *L*_*r*_ is computed on the reconstructed output from the decoder and used for pretraining the model. *L*_*s*_ and *L*_*pl*_ regulate the latent space for sparsity and semantic locality properties and make the representation more suitable for biclustering.

**Fig. 2.**
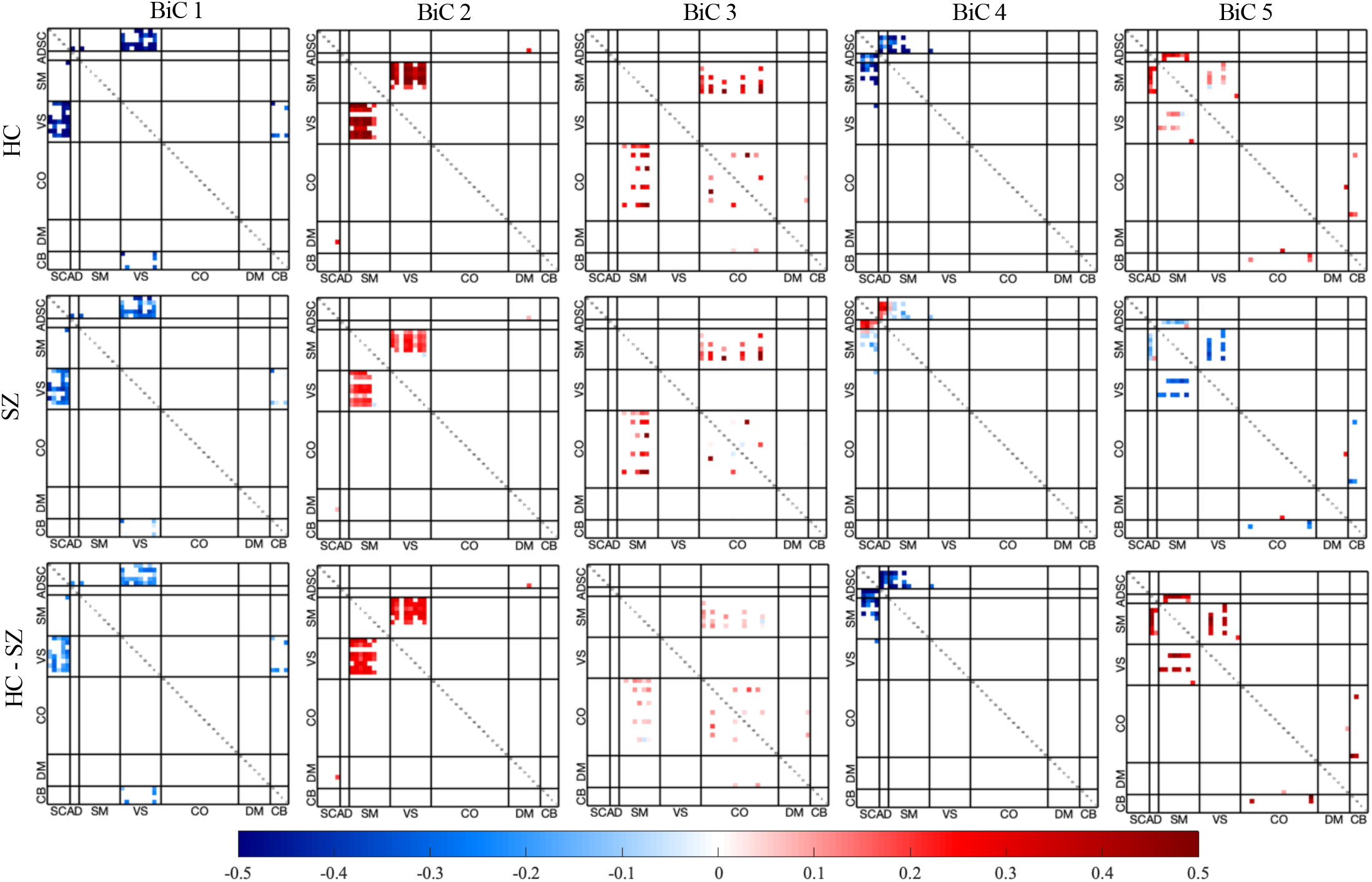
Static functional network connectivity (sFNC) patterns in the biclusters. The top row represents the sFNC matrices computed by averaging the HC subjects within a bicluster. The middle row for SZ and bottom HC-SZ difference. Each cell in these matrices represents the average connectivity strength for a given connection across the subject group included in a bicluster. The color bar represents the strength of the connection.

The highlights of our main contributions are as follows,

- A deep neural network (DNN) for biclustering
- Introduced semantic locality in features subgrouping for better neuro-interpretability.
- Sparsity regularization for improving representation learning and with enriched separability.
- Not bounded to discrete data. It can operate on continuous data representation learning and biclustering.

## II. Data Preprocessing And Static FNC (SFNC)

Our dataset is a combination of three studies, COBRE [11], fBIRN [12], and MPRC [13] consisting of 437 subjects with 275 healthy control (HC) and 162 schizophrenic (SZ) subjects.

The functional magnetic resonance imaging (fMRI) data is preprocessed by statistical parametric mapping (SPM12, http://www.fil.ion.ucl.ac.uk/spm/). The preprocessing steps include head motion correction and slice-timing correction for timing differences in slice acquisition. The detailed preprocessing steps are consistent with these studies [14, 15]. We run spatially constrained group ICA (gICA) from the NeuroMark [16] pipeline on the fMRI scans. gICA decomposes the imagery and finalizes 53 intrinsic connectivity networks (ICNs) with their time courses (TCs) which are grouped into seven brain domains [15]. The functional connection is usually estimated using the Pearson correlation between ICNs over time. The mean across the time course generates a 53 × 53 symmetric square matrix popularly known as static functional network connectivity (sFNC) [15].

## III. Semantic Locality Preserving Auto Decoder (SpaDE)

SpaDE is built on an autoencoder (AE) architecture equipped with specialized regularizations. It imposes two additional constraints on the latent space of the AE to make it more separable and cluster-friendly. After the model converges, we use a meta-heuristic (discussed next) on the hidden activation and weight matrix inspired by this study [17] to decide the bicluster assignments of features and samples. Fig. 1 illustrates the proposed SpaDE architecture. It incorporates two branches of neural networks known as an encoder for compression and a decoder as the mirror image of the encoder for decompression. Both the encoder and decoder consist of two linear layers followed by ReLu activation. The idea is to send the data through the bottleneck where the size of the bottleneck (i.e., the number of neurons in the hidden layer) is specified by the number of expected biclusters (*k*). The sFNC matrix is square and symmetric thus, we take the upper diagonal of the connectivity matrix and vectorize it for training purposes. We decide the most feasible configuration of the network including the number of biclusters *k* using the grid search evaluated on the reconstruction loss shown in (1). The other constraints we use on our latent representations are semantic locality (2) and sparsity constraints (4). Let the data matrix be *X*: *N × M, N* number of samples, and *M* features. Since our bicluster construction depends on the learned weight matrix (*W*) and hidden activation (*α*) of the neural network, the constraints are designed to regulate the weight matrix, in general. The reconstruction loss for pre-training the model is,

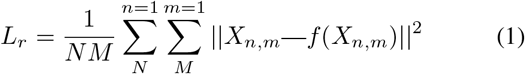

where *f* (*X*_*n,m*_) is the autoencoder’s reconstruction of the input *X*_*n,m*_ and ∥.∥ is the Euclidean norm. The constraint we use for preserving the semantic locality in the subgrouping is,

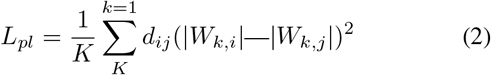

where *d*_*ij*_ is the similarity measure between two features *i, j* in the data matrix. The similarity measure (*d*_*ij*_) is defined as follows,

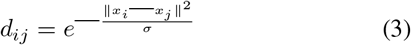

Equation (3) is inspired by the Gaussian kernel [31] where ∥*x*_*i*_—*x*_*j*_∥ is the distance between *x*_*i*_ and *x*_*j*_ and *σ* ∈ ℝ is a tuning parameter. We computed the distance using a special adaptation of earth mover distance (EMD) for one-dimensional vectors [18] hence suitable for our data (sFNC) - vectorized and bounded [—1 1]. Notice that for a smaller value of ∥*x*_*i*_—*x*_*j*_∥ and *σ*, the heuristic keeps the nearby data points closer. Then, we impose a sparsity constraint on the weight matrix to avoid the bias from larger weights eventually helps better reconstruct the features. We use a revised *L*1 norm on the *W* matrix for the sparsity. However, we also control for the sparsity in the primary decomposition of neuroimaging data using group ICA. Therefore, we down-scale the sparsity penalty in error calculation compared to the standard *L*1 norm.

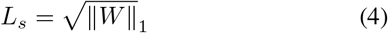

The objective function using these constraints is formulated as,

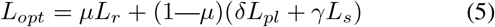

where *δ* and *γ* are non-negative and control the regularization by different penalty terms. *µ* helps train the model by balancing the load between feature learning and cluster-oriented loss. Through learning the reconstruction of the data matrix with these constraints, the model captures the contribution of each sample and feature characterized by the entries in the weight matrix and layer activation [17]. After the training scheme, the weight |*W*_*k,f*_ | determines the contribution of the *f*^*th*^ feature to the *k*^*th*^ bicluster, similarly, 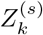 decides for *s*^*th*^ subject. The meta-heuristic for bicluster’s inclusion criteria is,

- Subject selection: Pick any subject *s* with 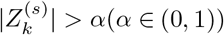
- Feature selection: Pick any feature *f* with |*W*_*k,f*_ | > *β*(*β* ∈ (0, 1))

### Model Training

The auto-encoder model is pre-trained using the reconstruction error described in (1) till achieving a moderate performance on recovering the input sequence. Then, we impose the clustering constraints on the latent space. For fine-tuning, we use the semantic locality and sparsity penalties described in (2) and (4) respectively. The overall optimization objective is defined in (5) and minimized using Adam optimizer with a learning rate of 10^−4^. In training, we start with a large value for *µ* that regulates the representation learning with reconstruction loss. Then, we reduce *µ* to enable the cluster-oriented influences on the latent space. The framework is built in Pytorch and trained for 450 epochs.

## IV. Results

We use several biclustering methods to compare SpaDE performance on three extensively researched datasets MPRC, fBIRN, and COBRE. We select two empirical biclustering methods FABIA [19] and N-BiC [2]. Then, we incorporate more concurrent deep learning models Auto Decoder (AD) [17], GAEBiC [20], and GraphMAE [21]. The functional connectivity is also modeled by using a graph neural network (GNN). For a more comprehensive comparison, we included GraphMAE and GAEBic. Moreover, we add two ablation studies to check the efficacy of the proposed constraints. Ablation 1 is without SLP and ablation 2 represents SpaDE without sparsity loss. We run the models for 100 repetitive iterations and present the *(mean ± standard deviation)* performance across the runs.

### Performance Metrics

We don’t have any ground truth partition for the combined dataset. So, we introduce a metric based on functional connectedness [22] to measure coherence in the bi-clustered community.

### Functional Coherence (FCO)

FCO compares the strength of the connectivity among the attributes within a bicluster and the interactions with the other parts of the system. In brain dynamics, it’s a measure of the interaction among a subset of brain networks and their communication with the rest of the brain. Given the sFNC matrices, to evaluate a partitioning *B*, we can measure the inter and intra connectedness (*C*) of a bicluster *b* as follows,

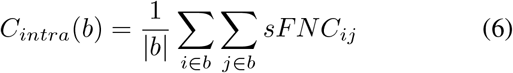

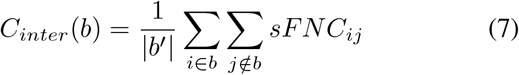

where *b*^*′*^ is the set of connections made by the community *b* with the rest of the system. We formulate functional coherence (FCO) for a biclustering run *B* as in the equation (8).

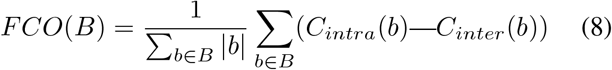

We also adapt two frequently used evaluation metrics mean square residue (MSR) [23] and average Pearson correlation coefficients (APCC) [24] for performance comparison. Table I shows the performance comparison between our model a baseline (*k*-means) and state-of-the-art biclustering methods. We run our experiments for three different *k* values that represent the number of biclusters. We obtain the best performance for *k* = 5. Our proposed framework outperforms various comparing methods by a margin in the performance metrics we included (Table I). The ablation experiments show slight degradation in performance which validate the efficacy of the constraints governing the biclustering process.

**TABLE I.**
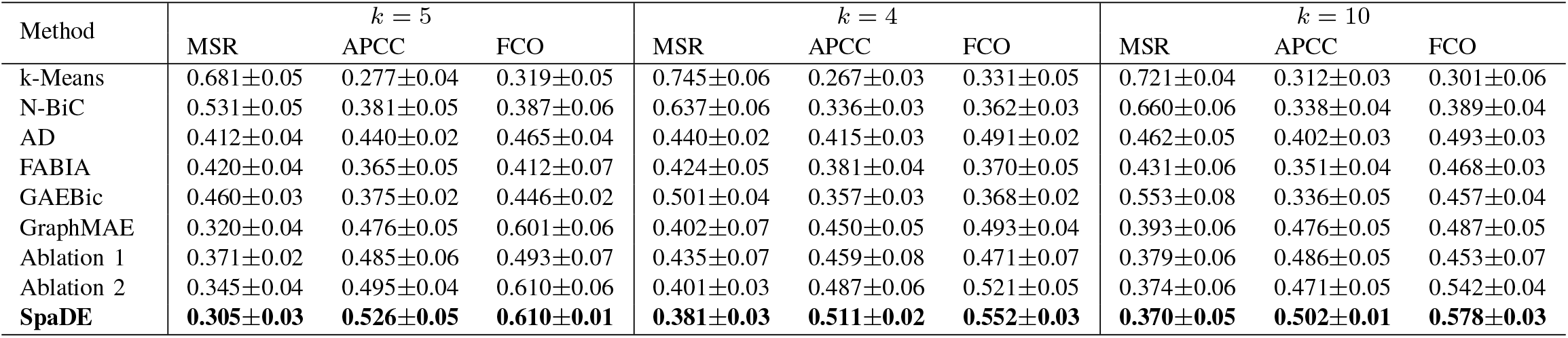
BICLUSTERING PERFORMANCE COMPARISON. THE HIGHLIGHTED BOTTOM ROW SHOWS THE BEST PERFORMANCES.

| Method | $k = 5$ | | | $k = 4$ | | | $k = 10$ | | |
| --- | --- | --- | --- | --- | --- | --- | --- | --- | --- |
|  | MSR | APCC | FCO | MSR | APCC | FCO | MSR | APCC | FCO |
| k-Means | 0.681±0.05 | 0.277±0.04 | 0.319±0.05 | 0.745±0.06 | 0.267±0.03 | 0.331±0.05 | 0.721±0.04 | 0.312±0.03 | 0.301±0.06 |
| N-BiC | 0.531±0.05 | 0.381±0.05 | 0.387±0.06 | 0.637±0.06 | 0.336±0.03 | 0.362±0.03 | 0.660±0.06 | 0.338±0.04 | 0.389±0.04 |
| AD | 0.412±0.04 | 0.440±0.02 | 0.465±0.04 | 0.440±0.02 | 0.415±0.03 | 0.491±0.02 | 0.462±0.05 | 0.402±0.03 | 0.493±0.03 |
| FABIA | 0.420±0.04 | 0.365±0.05 | 0.412±0.07 | 0.424±0.05 | 0.381±0.04 | 0.370±0.05 | 0.431±0.06 | 0.351±0.04 | 0.468±0.03 |
| GAEBic | 0.460±0.03 | 0.375±0.02 | 0.446±0.02 | 0.501±0.04 | 0.357±0.03 | 0.368±0.02 | 0.553±0.08 | 0.336±0.05 | 0.457±0.04 |
| GraphMAE | 0.320±0.04 | 0.476±0.05 | 0.601±0.06 | 0.402±0.07 | 0.450±0.05 | 0.493±0.04 | 0.393±0.06 | 0.476±0.05 | 0.487±0.05 |
| Ablation 1 | 0.371±0.02 | 0.485±0.06 | 0.493±0.07 | 0.435±0.07 | 0.459±0.08 | 0.471±0.07 | 0.379±0.06 | 0.486±0.05 | 0.453±0.07 |
| Ablation 2 | 0.345±0.04 | 0.495±0.04 | 0.610±0.06 | 0.401±0.03 | 0.487±0.06 | 0.521±0.05 | 0.374±0.06 | 0.471±0.05 | 0.542±0.04 |
| **SpaDE** | **0.305±0.03** | **0.526±0.05** | **0.610±0.01** | **0.381±0.03** | **0.511±0.02** | **0.552±0.03** | **0.370±0.05** | **0.502±0.01** | **0.578±0.03** |

### Biclusters Analysis

Figure 2 visualizes the sFNC connections for the identified biclusters. These localized patterns increase the chance of conducting uniform neuronal activity such as motor, visual, and auditory responses. It implies semantic localization through functional coherence among the connections in a bi-clustered subgroup. In bicluster 2, visual and sensorimotor connections are supportive of motor learning, and their impairment is acknowledged in schizophrenia [25]. Also, the auditory and subcortical connections in bicluster 1 might be contributing to the behavioral changes in schizophrenia [26]. Bicluster (BiCs) 1, 2, 4, and 5 show significant connectivity differences mostly among visual (VS), subcortical (SC), and sensorimotor (SM). Neuroimaging research has identified these domains to be associated with schizophrenia dysfunction and social impairments [25]. Reduced connectivity strength (BiC 2) and opposite connectivity (BiC 5) between SM and VS are conceivably responsible for the obstruction in social cognition and mentalization [27, 25]. The biclusters also depict significant group differences between schizophrenia (SZ) and the healthy control (HC) cohort. The connections in BiC 4 and 5 show reverse directionality in HC and SZ groups. It indicates the relevant cognition driven by these connected communities is divergent which may justify the cognitive differences between the subject groups. However, group differences in BiC 1, 2, and 3 are induced by the connectivity strength. In general, the SZ neural components are weakly connected compared to HC.

### Association with Cognitive Scores

Figure 3 shows the association of bi-clustered sFNC patterns with multiple cognitive tasks. The design of cognitive tests and acquisition of cognitive scores for the datasets are described in this study [28]. The studies used CMINDS [29] and MCCB [30] batteries to measure the cognitive scores. Additionally, we conduct standard data harmonization procedures to ensure consistency in scores across datasets. Our analysis reveals that BiC 1 exhibits notably higher correlations with all cognitive measures, underscoring the importance of interactions between SC and VS systems in task performance. This trend extends to BiC 5, which demonstrates connections between sensorimotor to auditory (AD) and VS systems, particularly relevant for auditory processing. Conversely, VS-SM connectivity in BiC 2 displays a strong negative correlation with problem-solving performance. That characterizes the relatedness of inverse directional flow for specific cognitive tasks. In summary, the results show sensorimotor and visual domains play crucial roles in a variety of motor and visual processing tasks [27].

**Fig. 3.**
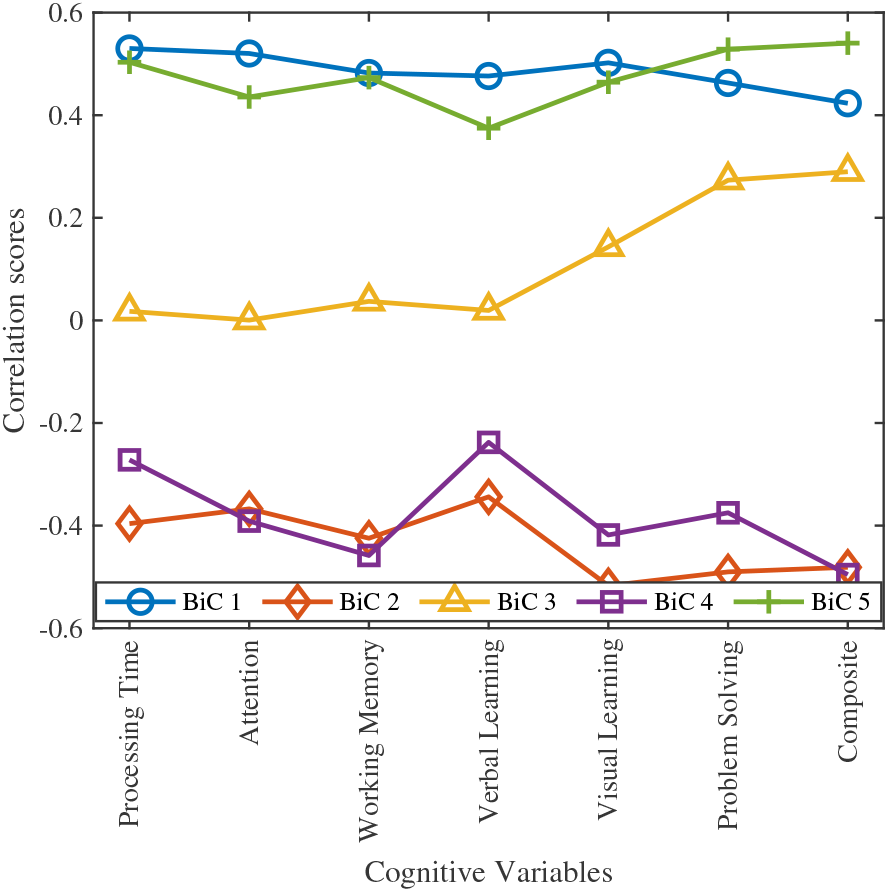
Correlation of bi-clustered sFNC patterns with distinct cognitive variables

## V. Conclusions

In this paper, we proposed a deep biclustering model for delving into smaller but meaningful nuances of neuroimaging data. The method aims to utilize the contextual knowledge in the data matrix for clustering and minimize overlap among the partitions. It reveals semantically cohesive and modular communities in the brain’s functional connectivity. The experiments demonstrate their association with cognitive performance and how the connectivity patterns differ in patients and healthy controls. The performance analysis shows that our method provides a better subgrouping of samples with a significant improvement in biclusters’ quality. Also, this framework is extendable to multi-modal data and multi-omics clusters which might provide inter-modality biclusters for exploring homogeneity across multiple physiological sources.

## Notes

### Competing Interest Statement

The authors have declared no competing interest.

